# *In vivo* multi-parametric manganese-enhanced MRI for detecting senile plaques in rodent models of Alzheimer’s disease

**DOI:** 10.1101/2021.01.12.426392

**Authors:** Eugene Kim, Davide Di Censo, Mattia Baraldo, Camilla Simmons, Ilaria Rosa, Karen Randall, Clive Ballard, Ben R Dickie, Steven CR Williams, Richard Killick, Diana Cash

## Abstract

Senile plaques are a hallmark of Alzheimer’s disease (AD) that develop in its earliest stages. Thus, non-invasive detection of these plaques would be invaluable for diagnosis and the development and monitoring of treatments, but this remains a challenge due to their small size. Here, we investigated the utility of manganese-enhanced MRI (MEMRI) for visualizing plaques in transgenic rodent models of AD across two species: 5xFAD mice and TgF344-AD rats.

Fourteen mice (eight transgenic, six wild-type) and eight rats (four transgenic, four wild-type) were given subcutaneous injections of MnCl_2_ and imaged *in vivo* using a 9.4T Bruker scanner. Susceptibility-weighted images, transverse relaxation rate (R2*) maps, and quantitative susceptibility maps were derived from high-resolution 3D multi-gradient-echo (MGE) data to directly visualize plaques. Longitudinal relaxation rate (R1) maps were derived from MP2RAGE data to measure regional manganese uptake. After scanning, the brains were processed for histology and stained for beta-amyloid (4G8 antibody), iron (Perl’s), and calcium/manganese (Alizarin Red).

MnCl_2_ improved signal-to-noise ratio (1.55±0.39-fold increase in MGE images) as expected, although this was not necessary for detection of plaques in the high-resolution images. Plaques were visible in susceptibility-weighted images, R2* maps, and quantitative susceptibility maps, with increased R2* and more positive magnetic susceptibility compared to surrounding tissue.

In the 5xFAD mice, most MR-visible plaques were in the hippocampus, though histology confirmed plaques in the cortex and thalamus as well. In the TgF344-AD rats, many more plaques were MR-visible throughout the hippocampus and cortex. Beta-amyloid and iron staining indicate that, in both models, MR visibility was driven by plaque size and iron load.

Voxel-wise comparison of R1 maps revealed increased manganese uptake in brain regions of high plaque burden in transgenic animals compared to their wild-type littermates. Interestingly, in contrast to plaque visibility in the high-resolution images, the increased manganese uptake was limited to the rhinencephalon in the TgF344-AD rats (family-wise error (FWE)-corrected p < 0.05) while it was most significantly increased in the cortex of the 5xFAD mice (FWE-corrected p < 0.3). Alizarin Red staining suggests that manganese bound to plaques in 5xFAD mice but not in TgF344-AD rats.

Multi-parametric MEMRI is a simple, viable method for detecting senile plaques in rodent models of AD. Manganese-induced signal enhancement can enable higher-resolution imaging, which is key to visualizing these small amyloid deposits. We also present *in vivo* evidence of manganese as a potential targeted contrast agent for imaging plaques in the 5xFAD model of AD.

**Highlights:**

- This is the first study to use manganese-enhanced MRI (MEMRI) for direct visualization of senile plaques in rodent models of Alzheimer’s disease, *in vivo*.
- Manganese enhancement is not necessary to detect plaques but improves image contrast and signal-to-noise ratio.
- Manganese binds to plaques in 5xFAD mice but not in TgF344-AD rats, demonstrating potential as a targeted contrast agent for imaging plaques in certain models of AD.

## 1 Introduction

Senile plaques (extracellular deposits of beta-amyloid (Aβ) in the brain) are one of the two neuropathological hallmarks of Alzheimer’s disease (AD) and develop early in the disease progression. However, early diagnosis of AD is limited by the difficulty of visualizing Aβ plaques *in* vivo. Currently, a definitive diagnosis of Alzheimer’s disease (AD) is only made *postmortem* by observing senile plaques and neurofibrillary tangles (intracellular deposits of hyperphosphorylated forms of the tau protein) in brain sections.

A diagnosis of AD during life can be greatly aided by positron emission tomography detection of radioactive Aβ ligands such as PiB (Mathis et al., 2012). Although highly accurate (Jack et al., 2018), the implementation of PET is restricted by high cost, limited accessibility, and invasiveness (ionizing radiation). Thus, non-invasive and repeatable methods of detecting plaques, or other facets of AD pathology, are needed to provide biomarkers of AD for refining diagnosis and assessing therapeutic efficacy.

MRI has shown great potential in filling this need (Ten Kate et al., 2018), but *in vivo* MR imaging of senile plaques remains challenging due to the small size of plaques and the relatively low sensitivity of MRI.

Previous preclinical studies on *in vivo* MR imaging of AD plaques involved complex pulse sequences (Jack et al., 2004) or administration of T1-shortening gadolinium-based contrast agents (GBCA) to increase the signal-to-noise ratio (SNR). GBCAs enable high-resolution imaging at reduced scan times, but complex procedures are required to deliver them to the brain parenchyma, e.g., stereotactic surgery for intracerebroventricular injection (Petiet et al., 2012) or the use of ultrasound and microbubbles to transiently open the blood-brain barrier (Santin et al., 2013).

Like Gd(III), manganese(II) is paramagnetic and shortens T1 relaxation times, but manganese-based contrast agents have lower relaxivities (i.e., produce a smaller decrease in T1 per unit of contrast agent concentration) than GBCAs (Brandt et al., 2019). Unlike GBCAs, MnCl_2_, a contrast agent commonly used for manganese-enhanced MRI (MEMRI), readily crosses the blood-brain barrier and, as a calcium analog, is taken up by neurons (Massaad & Pautler, 2011). Thus, in addition to enhancing SNR and neuroanatomical contrast, MEMRI can provide functional information. Accordingly, MEMRI has been used to probe AD-related disruption of neural activity and shown both increased (Fontaine et al., 2017; Tang et al., 2016) and decreased (Badea et al., 2019; Perez et al., 2013) Mn(II) uptake in different mouse models.

An overview of the use of MEMRI in neurodegenerative models is given in the recent review article by Saar and Koretsky (Saar & Koretsky, 2018).

In this study, we investigated the feasibility of using Mn(II) as a GBCA alternative to enhance image contrast and SNR to aid the *in vivo* visualization of senile plaques. We tested our MEMRI technique in two rodent models of AD:

1. The well-characterized 5xFAD transgenic mouse model of AD, which express human APP with the Swedish (K670N/M671L), Florida (I716V), and London (V717I) mutations and human PSEN1 with the M146L and L286V mutations and start developing Aβ plaques from as early as two months of age (Oakley et al., 2006).
2. The TgF344-AD transgenic rat model of AD, which express human APP with the Swedish mutation and human PSEN1 with the Δ exon 9 mutation and start developing Aβ plaques from as early as six months of age (Cohen et al., 2013).

## 2 Materials and methods

### 2.1 Experimental design

All experimental procedures involving animals were performed in accordance with the UK Animals (Scientific Procedures) Act 1986 and King’s College London institutional ethical guidelines.

All mice and rats were bred in in-house colonies. This study included eight 8–9.5-month-old 5xFAD mice, six of their wild-type littermates, four 16.5–18.5-month-old TgF344-AD rats, and four of their wild-type littermates (**Figure 1**). Half of the mice were male and half female across both genotypes, while all of the rats were male. A subset of six 5xFAD mice (3 male, 3 female) underwent baseline MRI scans. Immediately afterwards, these and all other animals received s.c. injections of MnCl_2_ (Sigma-Aldrich), once daily for four days. The mice received 0.15 mmol/kg/day (1.5 ml of 0.1M solution diluted in 1 ml of sterile 0.9% saline) for a cumulative dose of 0.6 mmol/kg; the rats received 0.075 mmol/kg/day (0.75 ml of 0.1M solution diluted in 1 ml of sterile 0.9% saline) for a cumulative dose of 0.3 mmol/kg. These doses were determined in a pilot study on a separate cohort of wild-type animals; a 0.6 mmol/kg cumulative dose resulted in mild adverse effects in the rats, thus a lower dose was used for this study.

**Figure 1.**
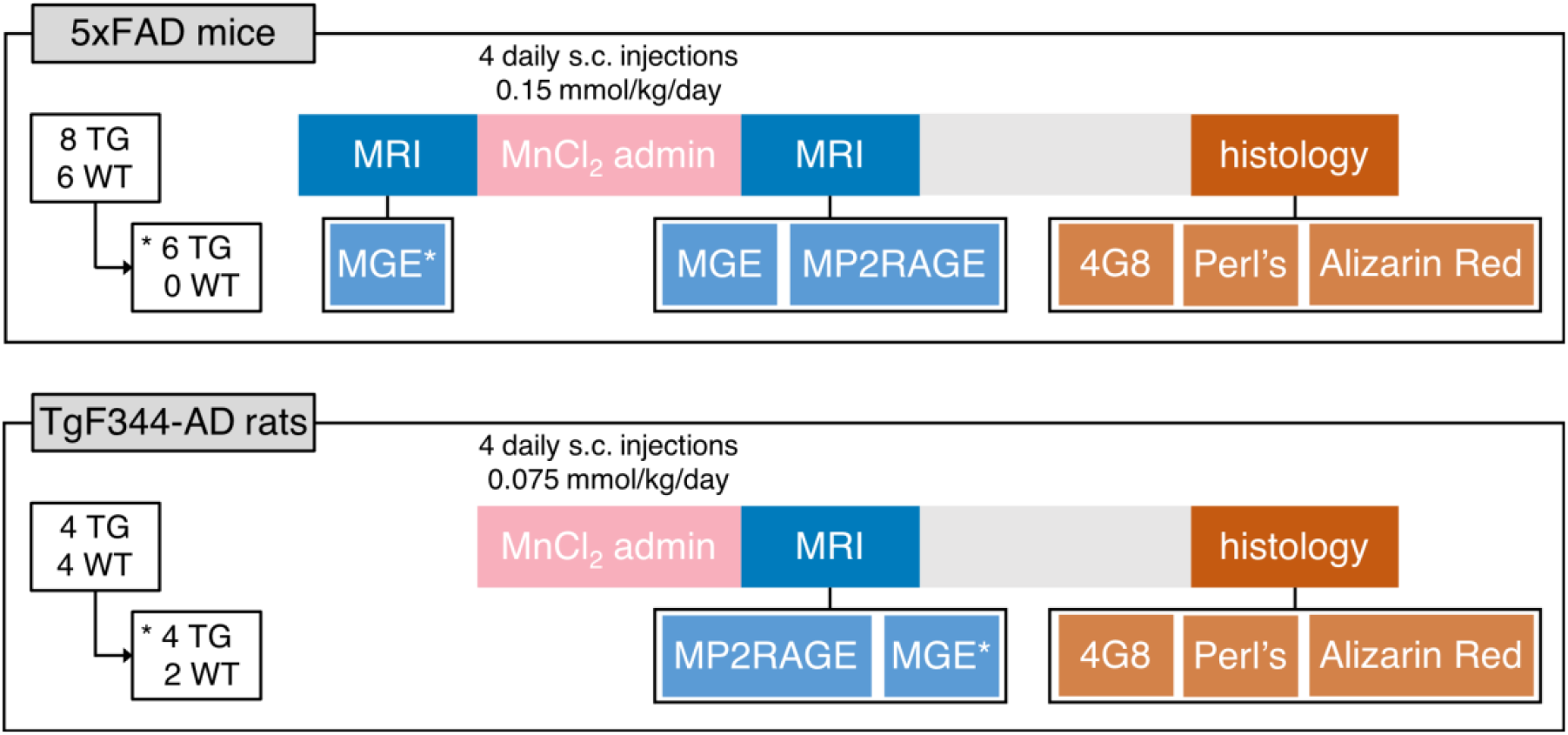
Experimental design. Fourteen mice (8 5xFAD transgenics [TG] and 6 wild-types [WT]) were scanned, including a subset of 6 TG mice that were scanned before MnCl_2_ administration (day 0). All mice were scanned after 4 days of daily s.c. MnCl_2_ injections (day 4). Eight rats (4 TgF344-AD TG and 4 WT) were scanned after MnCl_2_ administration (day 4). The MGE scan was only acquired for 4 TG and 2 WT rats. After the post-Mn scan, all animals were culled and their brains processed for histological analysis, which included 4G8 (Aβ), Perl’s (iron), and Alizarin Red (calcium/manganese) staining.

**Figure 2.**
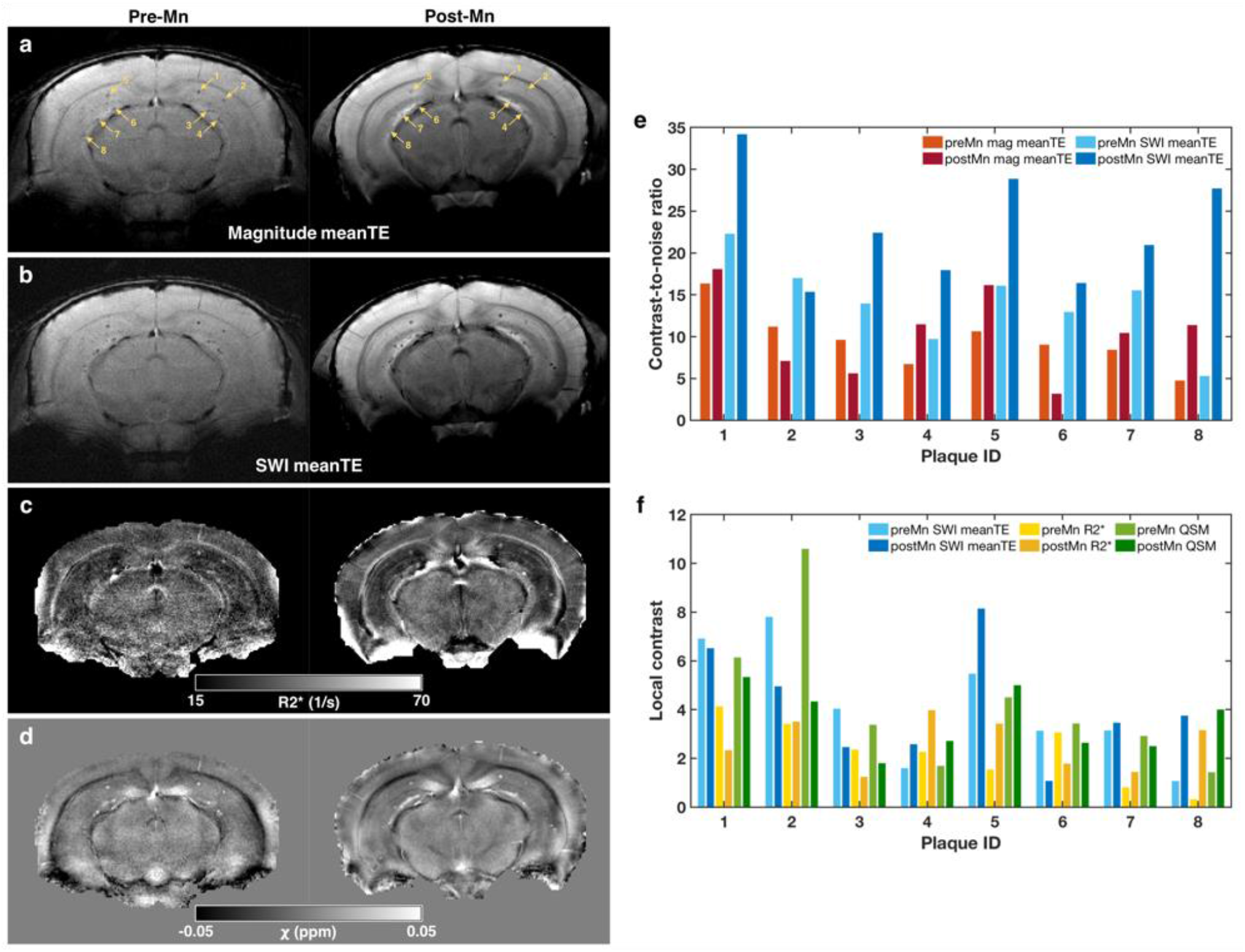
5xFAD plaque contrast in MGE-derived images and maps before and after MnCl_2_ administration. a) Matching slices of MGE images acquired from a 5xFAD mouse before (pre-Mn) and after (post-Mn) MnCl_2_. The images were computed by averaging the magnitude images across all four echo times (TE = 5-26 ms). Yellow arrows point to the same eight plaques, which were manually selected for contrast analysis. b-d) The same slices as in (a) but showing: b) the susceptibility-weighted images (SWI) averaged across TE, c) the R2* maps derived from mono-exponential fitting of the multi-echo data, and d) the magnetic susceptibility (X) maps derived from quantitative susceptibility mapping (QSM). e) Contrast-to-noise ratios (CNR) of each of the eight plaques in the pre- and post-Mn magnitude meanTE (a) and SWI meanTE images (b). CNR was calculated as abs(mean(plaque signal intensity) – mean(neighborhood signal intensity)) / sd(background signal intensity). f) Local contrast of each of the eight plaques in the pre- and post-Mn SWI meanTE images (b), R2* maps (c), and QSM maps (d). Local contrast was calculated as abs(mean(plaque signal intensity – mean(neighborhood signal intensity)) / sd(neighborhood signal intensity).

**Figure 3.**
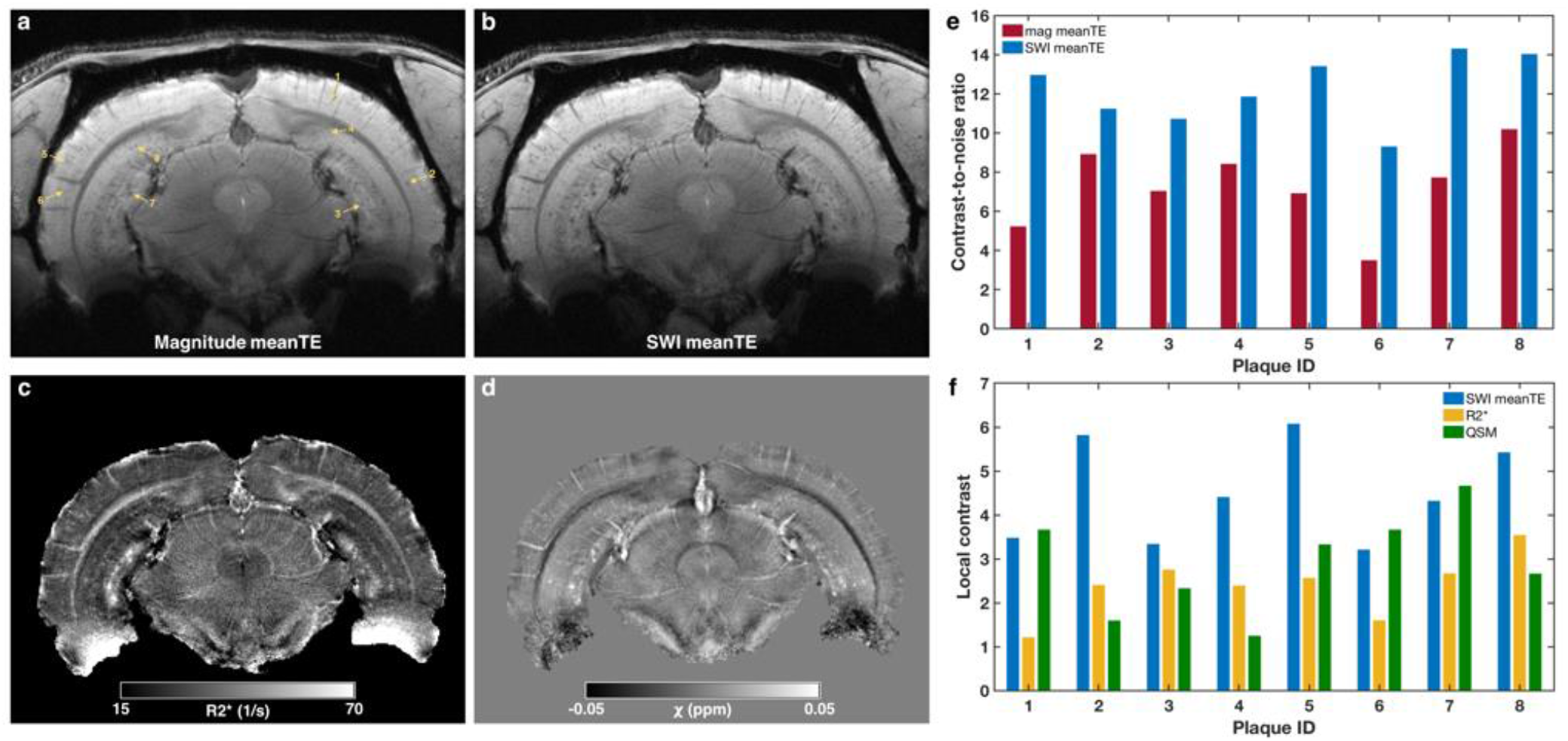
TgF344-AD plaque contrast in manganese-enhanced MGE-derived images and maps. a) A slice of an MGE image acquired from a TgF344-AD rat. The image was computed by averaging the magnitude images across all five echo times (TE = 6.2-40.8 ms). Yellow arrows point to eight plaques, which were manually selected for contrast analysis. b-d) The same slices as in (a) but showing: b) the susceptibility-weighted image (SWI) averaged across TE, c) the R2* map derived from mono-exponential fitting of the multi-echo data, and d) the magnetic susceptibility (X) map derived from quantitative susceptibility mapping (QSM). e) Contrast-to-noise ratios (CNR) of each of the eight plaques in the magnitude meanTE (a) and SWI meanTE images (b). CNR was calculated as abs(mean(plaque signal intensity) – mean(neighboring voxels signal intensity)) / sd(background signal intensity). f) Local contrast of each of the eight plaques in the SWI meanTE image (b), R2* map (c), and QSM map (d). Local contrast was calculated as abs(mean(plaque signal intensity – mean(neighboring voxels signal intensity)) / sd(neighboring voxels signal intensity).

MEMRI was performed on all animals one day after the final MnCl_2_ injection. Immediately after scanning, the animals were killed by transcardiac perfusion with heparinized saline and 4% formaldehyde. The fixed brains were harvested for histological analysis.

### 2.2 MRI acquisition

All MRI experiments were performed on a 9.4T Bruker BioSpec 94/20 controlled by ParaVision (6.0.1 for mice and 7.0.0 for rats) at the BRAIN Centre (http://brain-imaging.org) at King’s College London. An 86-mm volume coil was used in combination with species-specific, receive-only 2×2 surface array coils designed for mouse or rat brain imaging. The animals were anesthetized with isoflurane (5% induction, ∼2% maintenance) in medical air (1 L/min) + medical oxygen (0.4 L/min). During scanning, the isoflurane level was adjusted to maintain the respiration rate at 75-85 breaths/min for mice and 50-60 breaths/min for rats. Body temperature was monitored with a rectal probe and maintained at 37±0.5°C using a warm water circulation system (Small Animal Instruments, Inc., Stony Brook, NY, USA). Images were acquired using the following sequences (sequence parameters are summarized in **Table 1**):

**Table 1.**
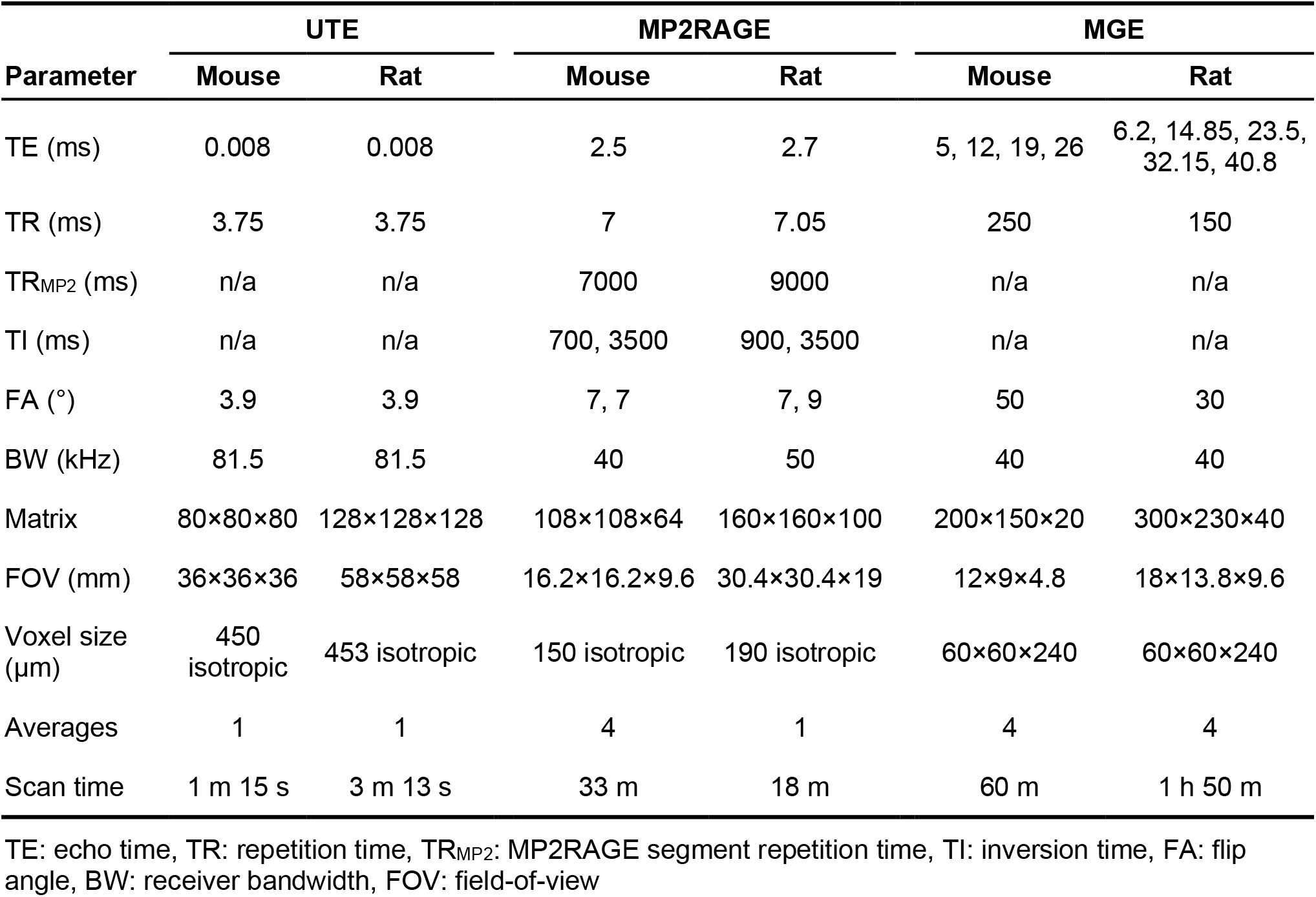
MRI sequence parameters

| Parameter | UTE |  | MP2RAGE |  | MGE |  |
| --- | --- | --- | --- | --- | --- | --- |
|  | Mouse | Rat | Mouse | Rat | Mouse | Rat |
| TE (ms) | 0.008 | 0.008 | 2.5 | 2.7 | 5, 12, 19, 26 | 6.2, 14.85, 23.5, 32.15, 40.8 |
| TR (ms) | 3.75 | 3.75 | 7 | 7.05 | 250 | 150 |
| TR<sub>MP2</sub> (ms) | n/a | n/a | 7000 | 9000 | n/a | n/a |
| TI (ms) | n/a | n/a | 700, 3500 | 900, 3500 | n/a | n/a |
| FA (°) | 3.9 | 3.9 | 7, 7 | 7, 9 | 50 | 30 |
| BW (kHz) | 81.5 | 81.5 | 40 | 50 | 40 | 40 |
| Matrix | 80×80×80 | 128×128×128 | 108×108×64 | 160×160×100 | 200×150×20 | 300×230×40 |
| FOV (mm) | 36×36×36 | 58×58×58 | 16.2×16.2×9.6 | 30.4×30.4×19 | 12×9×4.8 | 18×13.8×9.6 |
| Voxel size (μm) | 450 isotropic | 453 isotropic | 150 isotropic | 190 isotropic | 60×60×240 | 60×60×240 |
| Averages | 1 | 1 | 4 | 1 | 4 | 4 |
| Scan time | 1 m 15 s | 3 m 13 s | 33 m | 18 m | 60 m | 1 h 50 m |

TE: echo time, TR: repetition time, TR<sub>MP2</sub>: MP2RAGE segment repetition time, TI: inversion time, FA: flip angle, BW: receiver bandwidth, FOV: field-of-view

1. 3D ultra-short echo time (UTE)
2. Magnetization prepared 2 rapid acquisition of gradient echoes (MP2RAGE)
3. High-resolution 3D multi-gradient-echo (MGE)

The UTE and MP2RAGE sequences provided whole brain coverage, while the MGE sequence provided partial coverage along the rostrocaudal axis; the MGE slice package was positioned to include all of the hippocampus. The pre-Mn baseline scan sessions for the subset of six 5xFAD mice included only the MGE and UTE scans. All three scans were acquired in the post-Mn MEMRI sessions for all animals except for two wild-type rats, for which MGE scans were not acquired. One wild-type mouse was found to have hydrocephalus (**Figure 4**, second row, right) and thus excluded from the MP2RAGE processing and analysis described below.

**Figure 4.**
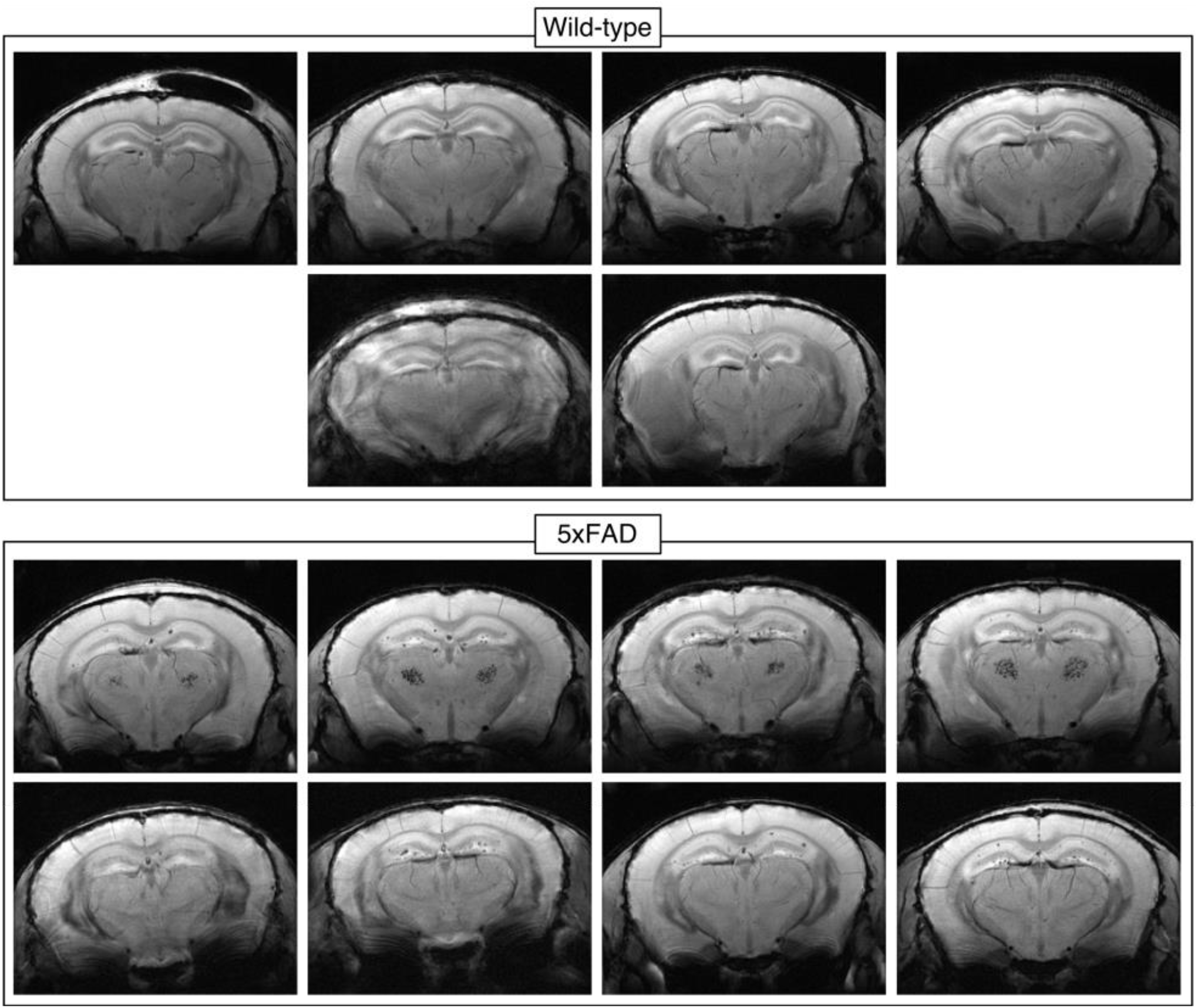
Plaques visible in the hippocampus of all 5xFAD mice. Single slices from MGE susceptibility-weighted meanTE images of all wild-type mice (one had severe Moiré fringe artifacts [second row, left], and one mouse had hydrocephalus [second row, right]) and all 5xFAD mice. Very few plaques were visible in one 5xFAD mouse due to Moiré fringe artifacts (bottom left). Half of the 5xFAD mice presented with bilateral hypointense clusters in the thalamus (third row) not found in wild-type littermates.

### 2.3 MRI processing and analysis

#### 2.3.1 MP2RAGE

Both magnitude and complex MP2RAGE and UTE images were reconstructed in ParaVision. The complex images from the four channels of the array coil were combined with the COMPOSER method, which uses the UTE reference scan to correct for differing phase offsets in the images from the individual coils in the array (Robinson et al., 2017). This was implemented in the wrapper script `composer.sh`, which is part of the QUantitative Imaging Tools (QUIT) package (https://github.com/spinicist/QUIT) (Wood, 2018). From the combined complex MP2RAGE image, bias-field-corrected T1w images and T1 maps were computed using the `mp2rage` command in QUIT, which produces robust T1w images by suppressing background noise (O’Brien et al., 2014). The noise suppression constant β was empirically optimized and set to 1 for all mouse data and 0.05 for all rat data. R1 maps were made by calculating the reciprocal of the T1 maps.

Study-specific T1w mouse and rat templates were created using the ANTs script `antsMultivariateTemplateConstruction2.sh`, and each subject was registered to its respective template using `antsRegistration` with serial rigid-body, affine, and SyN transformations (Avants et al., 2008).

Each R1 map was transformed to the template space, and R1 values were normalized by the median R1 value in a temporalis region-of-interest (ROI) manually defined on the template image. The rationale was to correct for any inter-subject differences in effective Mn(II) dose on the assumption that genotype did not affect Mn(II) uptake in the temporalis. To determine if R1 differed between transgenic and wild-type animals, voxel-wise permutation tests were performed using FSL `randomise` with 5000 permutations, threshold-free cluster enhancement, and controlling for family-wise error (FWE) rate (Winkler et al., 2014).

#### 2.3.2 MGE

Magnitude, complex, and susceptibility-weighted images (SWI) were reconstructed from the MGE data in ParaVision. The ‘positive-mask’ SWI reconstruction weighting mode was used, with a mask weighting of 4.0 and Gauss broadening of 0.2 mm.

The magnitude and susceptibility-weighted images were bias field corrected using the `N4BiasFieldCorrection` command in ANTs (Tustison et al., 2010). Then, individual echo time (TE) images were averaged to create magnitude and SWI “meanTE” images to increase SNR (Helms & Dechent, 2009). R2* maps were computed from the uncorrected MGE magnitude images using the non-linear fitting algorithm of the `multiecho` command in QUIT.

The complex images were combined as described above, from which magnitude and phase images were extracted for quantitative susceptibility mapping (QSM). Magnetic susceptibility (χ) maps were computed using the STAR-QSM algorithm (Wei et al., 2015) in STI Suite v3.0, a MATLAB (MathWorks, Natick, MA, USA) toolbox. Brain masks, which are required by the QSM algorithm, were generated from the uncorrected first TE magnitude images using the Rapid Automatic Tissue Segmentation (RATS) tool (Oguz et al., 2014).

Thus, four image contrasts or parametric maps were derived from the MGE data:

1. magnitude meanTE images,
2. SWI meanTE images,
3. R2* maps, and
4. QSM maps.

Using Fiji (Schindelin et al., 2012), the SNR of magnitude meanTE images were estimated from manually drawn ROIs around the brain and background in a central slice, where

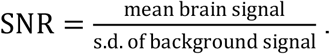

For one 5xFAD mouse and one TgF344-AD rat each, eight randomly selected plaques and their neighborhoods were manually segmented from a single slice of the SWI meanTE image using Fiji. Any voxels containing blood vessels or white matter were excluded from the neighborhoods. The contrast-to-noise ratios (CNR) in the magnitude and SWI meanTE images were computed for each plaque, where

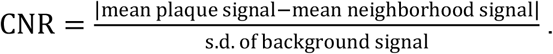

In addition, local contrast, defined here as

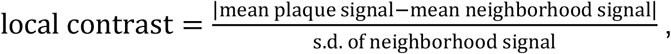

was calculated for each plaque in each of the four MGE-derived images/maps.

## 2.4 Histology

Perfusion-fixed brains were extracted and cryoprotected in 30% sucrose before being sectioned at 35 µm in a series of 6 for mouse and 12 for rat on a freezing microtome and stored free floating in cryoprotectant at −20°C. For mice and rats, Alizarin Red was used to stain for calcium and manganese, Perl’s/DAB for iron, and 4G8 antibody for Aβ.

For Alizarin Red, one series was mounted onto slides and air dried before rehydrating in distilled H_2_O and incubating in Alizarin Red solution for 2 min. Sections were then differentiated in acetone then acetone:xylene (1:1) and finally were cleared in xylene before coverslipping.

For Perl’s/DAB, one series was washed for 3 x 5 min in phosphate-buffered saline (PBS) before being incubated in 0.3% hydrogen peroxide (H_2_O_2_) for 30 min, then in Perl’s solution (1% Potassium Ferrocyanide Trihydrate in acidified PBS) for 1 hour at 37°C, and finally in 3,3′-Diaminobenzidine (DAB) for up to 10 min until sufficient colour had developed (PBS wash steps were performed in between each incubation). Sections were then mounted onto slides, air dried, and coverslipped.

For 4G8, one series was washed for 3 × 5 min in Tris-buffered saline (TBS) before being incubated in 88% formic acid for 15mins for antigen retrieval, then in 1% H_2_O_2_ for 15 min to block endogenous peroxidase activity, 10% skimmed milk powder to block non-specific binding, and finally in anti-4G8 antibody (1:2,000; BioLegend (800701)) overnight at 4°C. This was followed by incubation in a biotinylated secondary antibody (anti mouse in goat, 1:1000, Vector Labs (BA-9200)) for 2 hours and ABC kit for 1 hour (Vectastain ABC Kit, Vector Labs (PK-6100)). Washes with TBS-X (3 × 5 min) were performed in between each step. Staining was then visualized using DAB. Sections were then mounted onto slides, air dried, and coverslipped.

Slides from all three stains were then scanned with an Olympus VS120 slide scanner at ×40 magnification. Images were saved with 80% compression.

Raw MRI data is available on OpenNeuro (doi:10.18112/openneuro.ds003463.v1.0.0).

## 3 Results

### 3.2 Mn(II) increased the SNR of MGE images

One aim of this study was to evaluate the efficacy of using Mn(II) as a T1 shortening agent to increase the SNR and/or decrease the scan time of high-resolution images. To this end, high-resolution MGE images were acquired both before and after MnCl_2_ administration for six 5xFAD mice. The post-Mn SNR was on average 55% higher than the pre-Mn SNR, but the change varied greatly from 7% to 108% (**Table 2**). This variability may have been due to Mn(II)-independent variability in image quality stemming from sensitivities to shimming and motion of the long, gradient-echo-based scan.

**Table 2.**
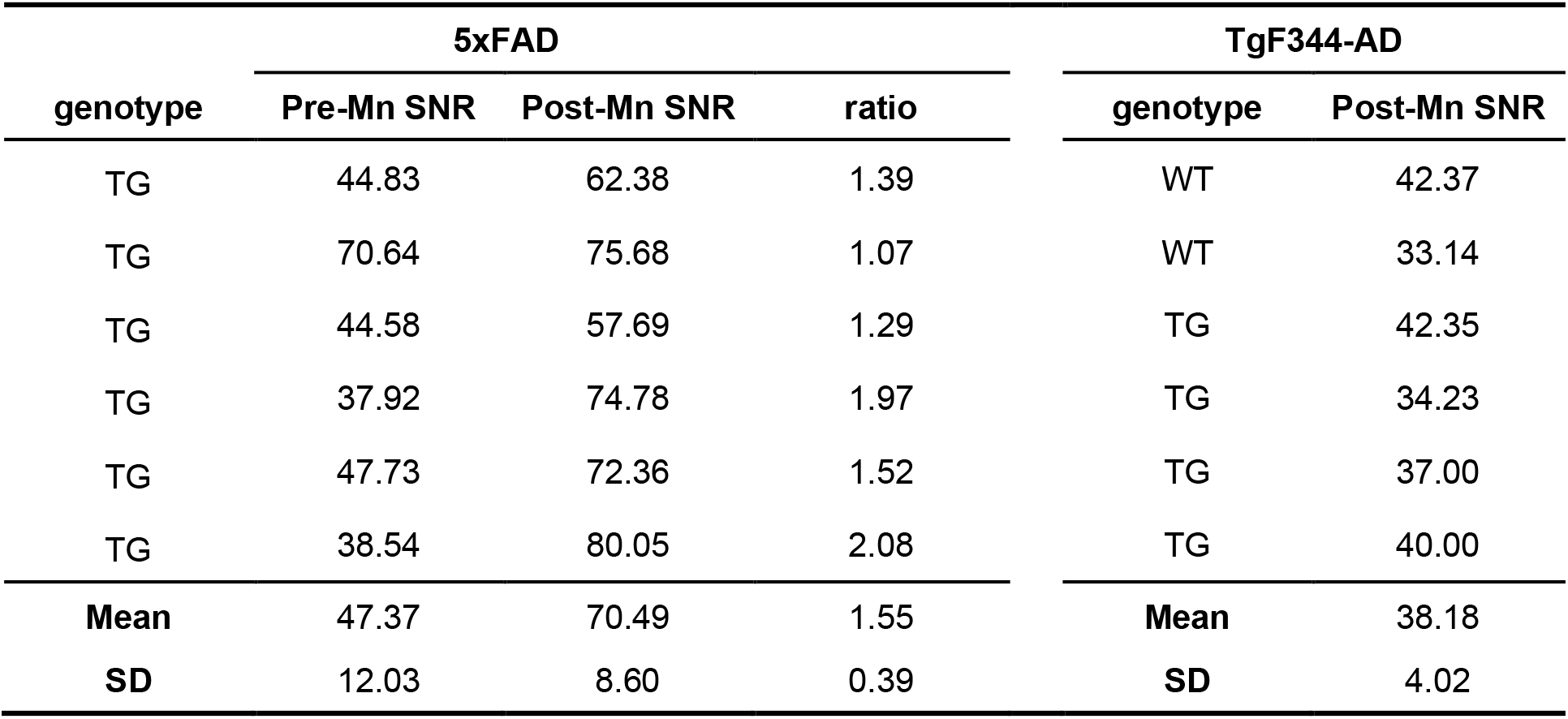
Signal-to-noise ratios of MGE magnitude meanTE images

Beyond increasing SNR, Mn(II) increased neuroanatomical contrast, most likely due to regional differences in uptake driven by neural activity and neuronal density (**Figure 2a**). This is apparent in the increased white/gray matter contrast and particularly in the visibility of hippocampal structures. This is far from a novel observation – Mn(II) is well-known to enhance neuroanatomical contrast – but it bears mentioning for this specific application because nominally T1-weighted FLASH-like sequences like the MGE sequence used in this study produce fairly flat image contrast at high fields. Thus, MEMRI was especially beneficial and aided more precise localization of plaques.

### 3.2 Mn(II) had a variable effect on plaque contrast

While Mn(II) increased global tissue contrast and thereby improved our ability to determine where plaques were located, it had a variable effect on our ability to detect plaques in the first place. Hypointense plaques were visible in magnitude and SWI meanTE images (**Figure 2a**,**b**) both before (left column) and after (right column) MnCl_2_ administration. Due to imperfect slice alignment between pre- and post-Mn scans, not all plaques visible in one were visible in the other. For one 5xFAD mouse, eight plaques visible in both pre- and post-Mn scans were randomly selected and manually segmented (yellow arrows, **Figure 2a**,**b**). The CNR of those plaques are presented in **Figure 2e**. In the magnitude meanTE images, the post-Mn CNR of three plaques were actually lower than the pre-Mn CNR; while in the SWI meanTE images, the post-Mn CNR was higher in all but one plaque. As mentioned above, the pre- and post-Mn slices were not perfectly aligned, which led to varying degrees of partial voluming in the slice direction and may have contributed to the unexpectedly lower post-Mn CNR of some plaques.

### 3.3 Plaque MR-visibility was driven by increased magnetic susceptibility

As for the 5xFAD mouse, we manually segmented eight random plaques in one TgF344-AD rat (**Figure 3**). All selected plaques in both the 5xFAD mouse and TgF344-AD rat had higher CNR in the SWI meanTE image compared to the magnitude image (**Figure 2e** and **Figure 3e**): 5xFAD post-Mn SWI CNR = 22.99 ± 6.70, magnitude CNR = 10.43 ± 5.07; TgF344-AD post-Mn SWI CNR = 12.23 ± 1.75, magnitude CNR = 7.25 ± 2.12. In addition, MGE-visible plaques had elevated R2* (**Figure 2c** and **Figure 3c**), which increases in the presence of magnetic susceptibility gradients. QSM confirmed that the plaques had a positive magnetic susceptibility greater than most of the surrounding brain parenchyma (**Figure 2d** and **Figure 3d**). Comparing the three image modalities – SWI, R2*, and QSM – the post-Mn local plaque contrast was generally highest in the SWI meanTE images (5xFAD: 4.12 ± 2.32, TgF344-AD: 4.51 ± 1.14), intermediate in the QSM maps (5xFAD: 3.54 ± 1.30, TgF344-AD: 2.90 ± 1.15), and lowest in the R2* maps (5xFAD: 2.61 ± 1.04, TgF344-AD: 2.40 ± 0.71). However, the relative local contrasts of the three modalities varied from plaque to plaque (**Figure 2f** and **Figure 3f**).

### 3.4 Only a fraction of histologically identified plaques were MR-visible

**Figure 4** and **Figure 5** show single slices of SWI meanTE images of each mouse and rat brain, respectively, for which MGE scans were acquired. Most visible plaques in 5xFAD mouse brains were in the hippocampus, while more plaques in TgF344-AD rat brains were visible in the cortex a well as the hippocampus. As expected, there were no visible plaques in any of the wild-type animals. Half of the 5xFAD mice presented with bilateral hypointense clusters in the thalamus, which were also not found in wild-type animals.

**Figure 5.**
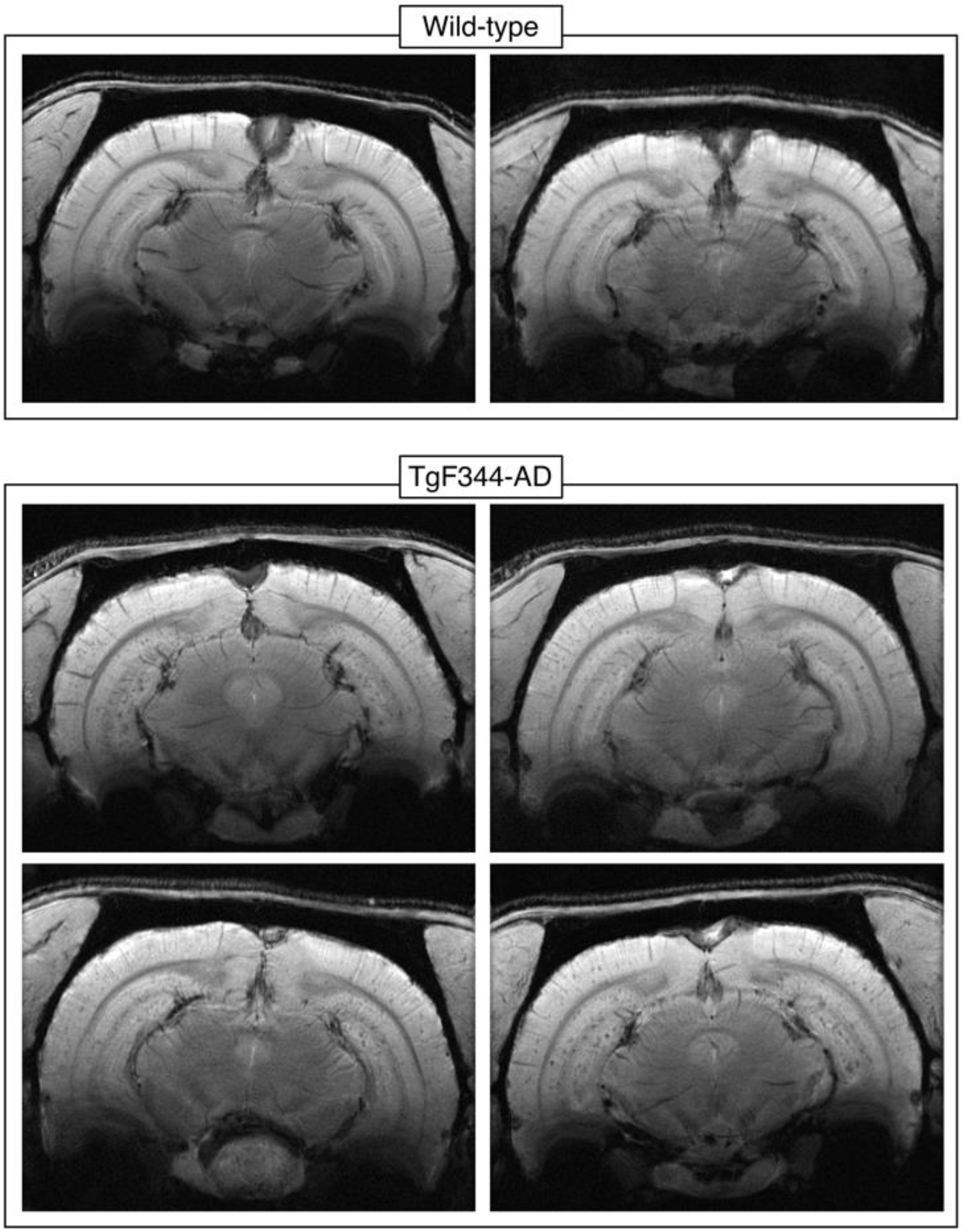
Plaques visible in the cortex and hippocampus of all TgF344-AD rats. Single slices from MGE susceptibility-weighted meanTE images of all wild-type rats and all TgF344-AD rats. Compared to the 5xFAD mice, the TgF344-AD plaques had lower contrast and were more obscured by blood vessels; still many more plaques were visible, especially in the cortex and ventral hippocampus.

4G8 anti-Aβ staining revealed senile plaques throughout the 5xFAD brain, with large plaque burdens in the septum, thalamus, and deeper cortical layers in addition to the hippocampus (**Figure 6a**). Perl’s/DAB staining showed that iron was present in many plaques in the cortex, hippocampus, and thalamus; but interestingly, very few plaques outside the hippocampus were visible in MGE images (**Figure 6b-d**).

**Figure 6.**
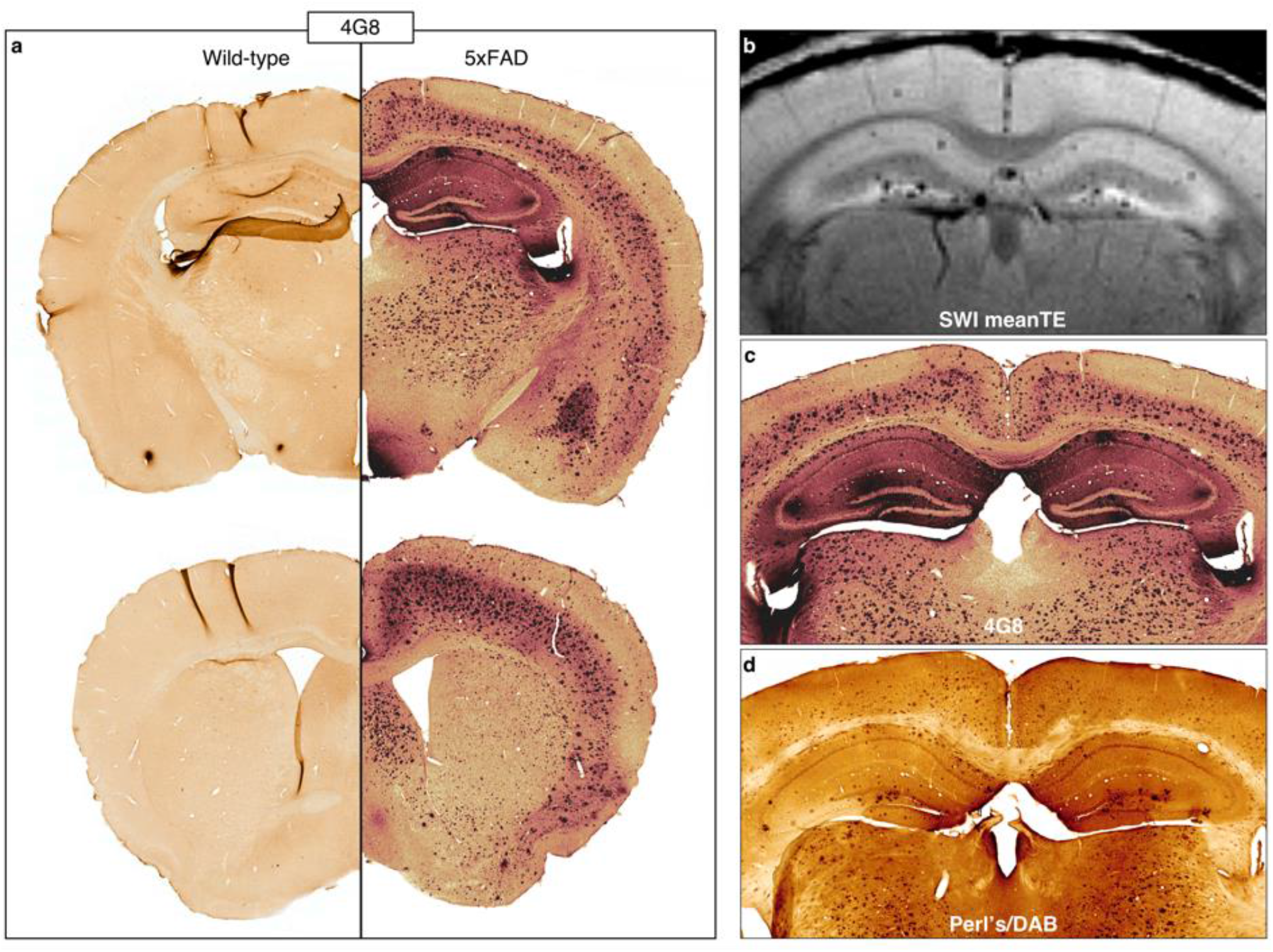
Histology shows iron-containing Aβ plaques throughout the brain in 5xFAD mice. a) 48G anti-amyloid staining confirmed the presence of plaques not only in the hippocampus, but in the deep cortical layers and subcortical regions, including the thalamus, of 5xFAD mice. b) The plaques visible on MRI were mostly limited to the hippocampus, with a few cortical plaques visible in a few mice. The contrast in these MGE images is largely T2*/susceptibility-driven. Perl’s-DAB staining showed that many of the plaques stained with 4G8 contained iron (c-d). Thus, iron load did not appear to play a large role in plaque MRI-visibility. Images in b-c were acquired from the same 5xFAD mouse.

The thalamic plaques in the 4G8- and Perl’s/DAB-stained sections (**Figure 6**) were not localized in the manner of the hypointense clusters seen in the MR images (**Figure 4**). QSM revealed that these clusters were diamagnetic at baseline (pre-Mn, **Figure 7a**) but became paramagnetic after MnCl_2_ administration (post-Mn, **Figure 7b**). Alizarin Red staining showed large spots in the same area of the thalamus (**Figure 7c**); these large spots were only present in mice with MR-visible thalamic clusters (**Figure 7d**).

**Figure 7.**
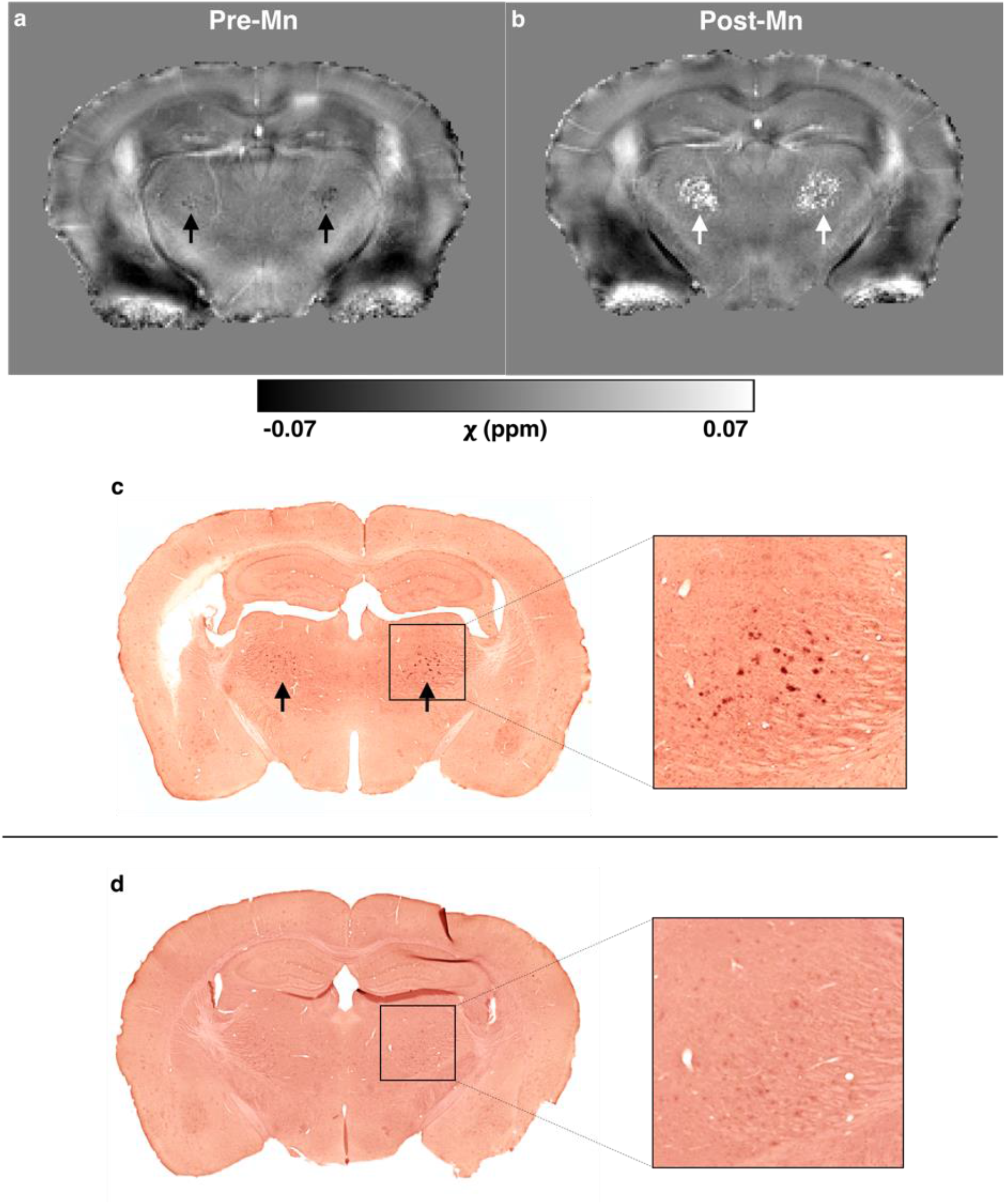
*In vivo* QSM and Alizarin Red staining reveal thalamic calcifications in 5xFAD mice. a-b) Quantitative susceptibility (χ) maps of a 5xFAD mouse a) before (pre-Mn) and b) after administration of MnCl_2_ (Post-Mn). Bilateral clusters in the thalamus were diamagnetic Pre-Mn but became paramagnetic Post-Mn (arrows). c) A section from the same mouse, stained with Alizarin Red, shows that these thalamic clusters contain calcium and/or manganese. d) An Alizarin-Red-stained section from a 5xFAD mouse without MR-visible thalamic clusters.

Alizarin Red staining also revealed numerous, less intensely stained foci in the brains of 5xFAD mice treated with MnCl_2_ (Mn+) that were not present in Mn+ wild-type brains (**Figures 7** and **8**). Fewer of these deposits were visible in the MnCl_2_-naïve (Mn-) 5xFAD brain, while the Mn- and Mn+ wild-type brain sections had a similar appearance. The Alizarin-Red-stained deposits in the Mn+ 5xFAD sections matched the spatial distribution of 4G8-stained Aβ plaques (**Figure 6a**), suggesting that the injected Mn(II) bound to the plaques. Unlike in the 5xFAD brains, Alizarin Red staining showed no evidence of Mn(II) binding to plaques in the TgF344-AD brains (**Figure 9**).

**Figure 8.**
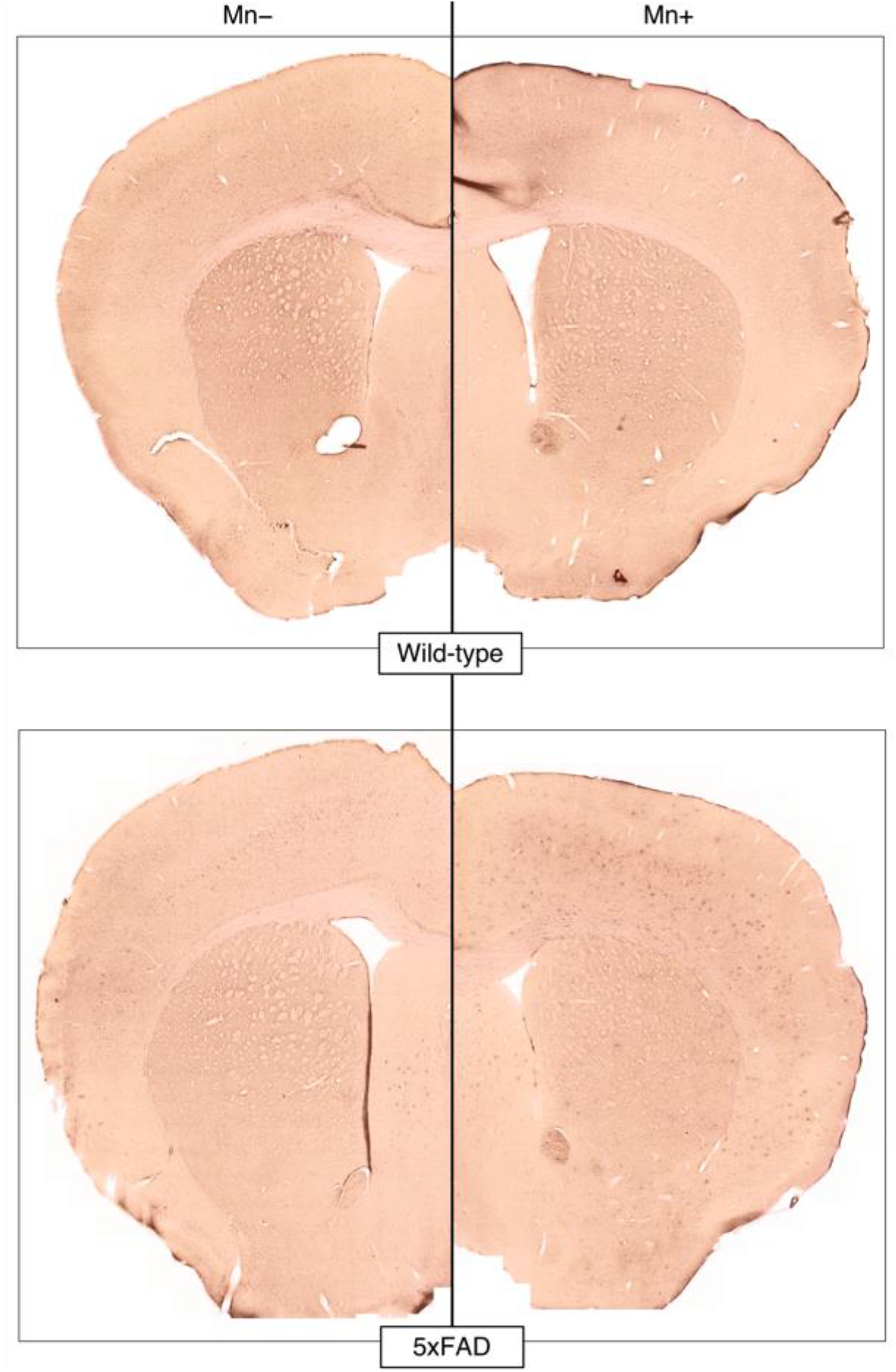
Alizarin Red staining shows subcutaneously administered manganese binds to Aβ plaques in the 5xFAD brain. Alizarin Red binds to calcium and, because of their chemical similarity, manganese. Alizarin Red staining appears similar in wild-type brains that were MnCl_2_-naïve (Mn-) and treated with MnCl_2_ (Mn+). In the Mn-5xFAD brain, some plaques or plaque-like structures in the cortex and septum were stained. Many more plaques were stained in the Mn+ 5xFAD brain, suggesting that the injected manganese bound to the plaques.

**Figure 9.**
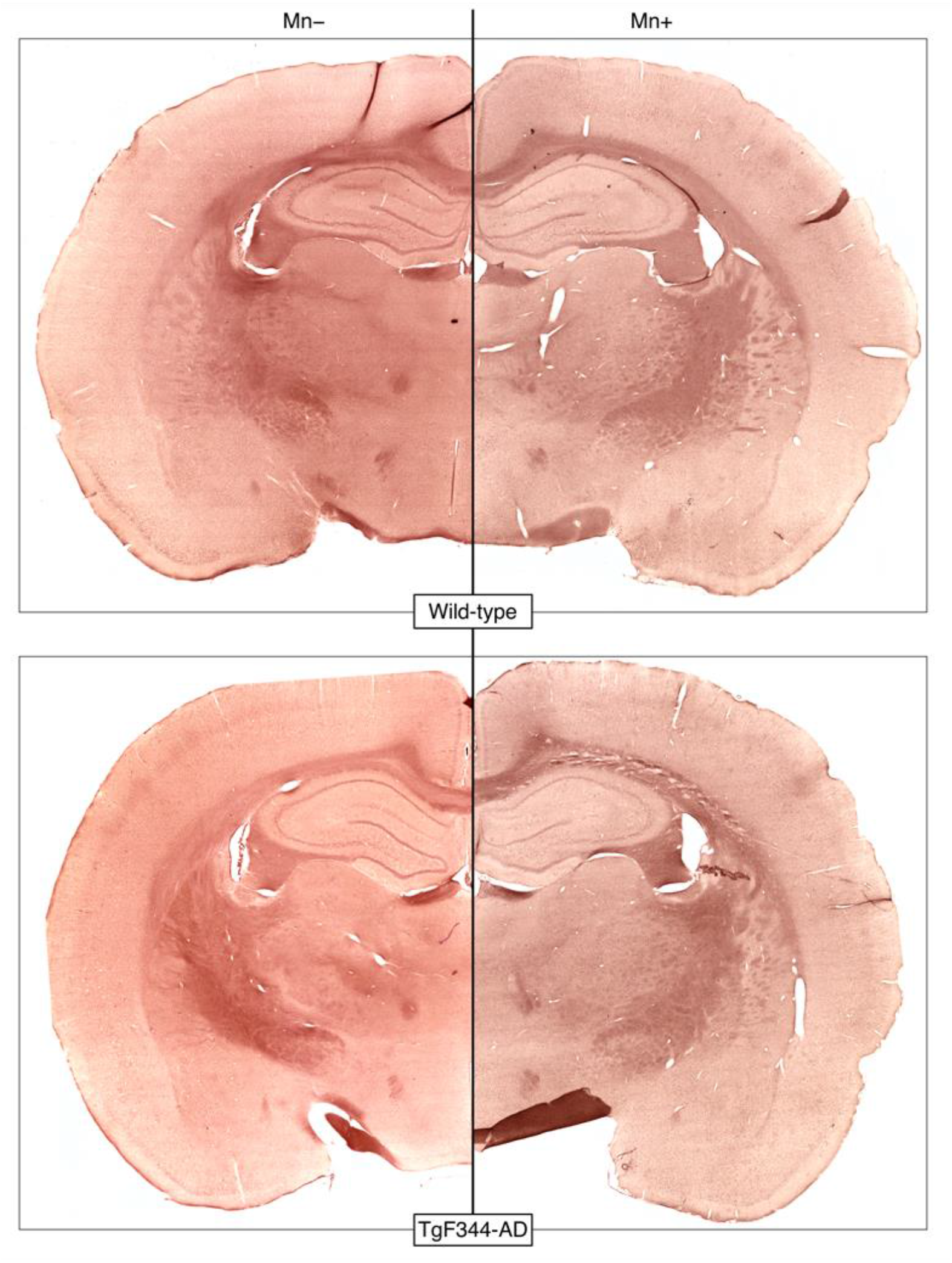
Alizarin Red staining reveals little difference between wild-type and TgF344-AD rat brains. Alizarin Red binds to calcium and, because of their chemical similarity, manganese. Alizarin Red staining appears similar in wild-type and TgF344-AD brains that were manganese-naïve (Mn-) and treated with MnCl_2_ (Mn+). Unlike in the 5xFAD mouse brains, there is no evidence that exogenously administered manganese bound to senile plaques in TgF344-AD rat brains.

Compared to 5xFAD mice, TgF344-AD rats had a lower plaque burden in general except in the hippocampus, amygdala, and piriform cortex (**Figure 10a**). Most plaques throughout the rat brain appeared to contain iron deposits (**Figure 10c**). The 4G8 and Perl’s/DAB staining revealed no obvious difference between plaques in the dorsal hippocampus versus those in the ventral hippocampus, but more plaques were MR-visible in the latter than in the former (**Figure 10b**). Plaques in the amygdalopiriform cortex were difficult to see in the MR images due to a combination of the receiver coil’s inhomogeneous sensitivity profile and the signal dropout around the air-tissue interfaces along the ventral surface of the brain.

**Figure 10.**
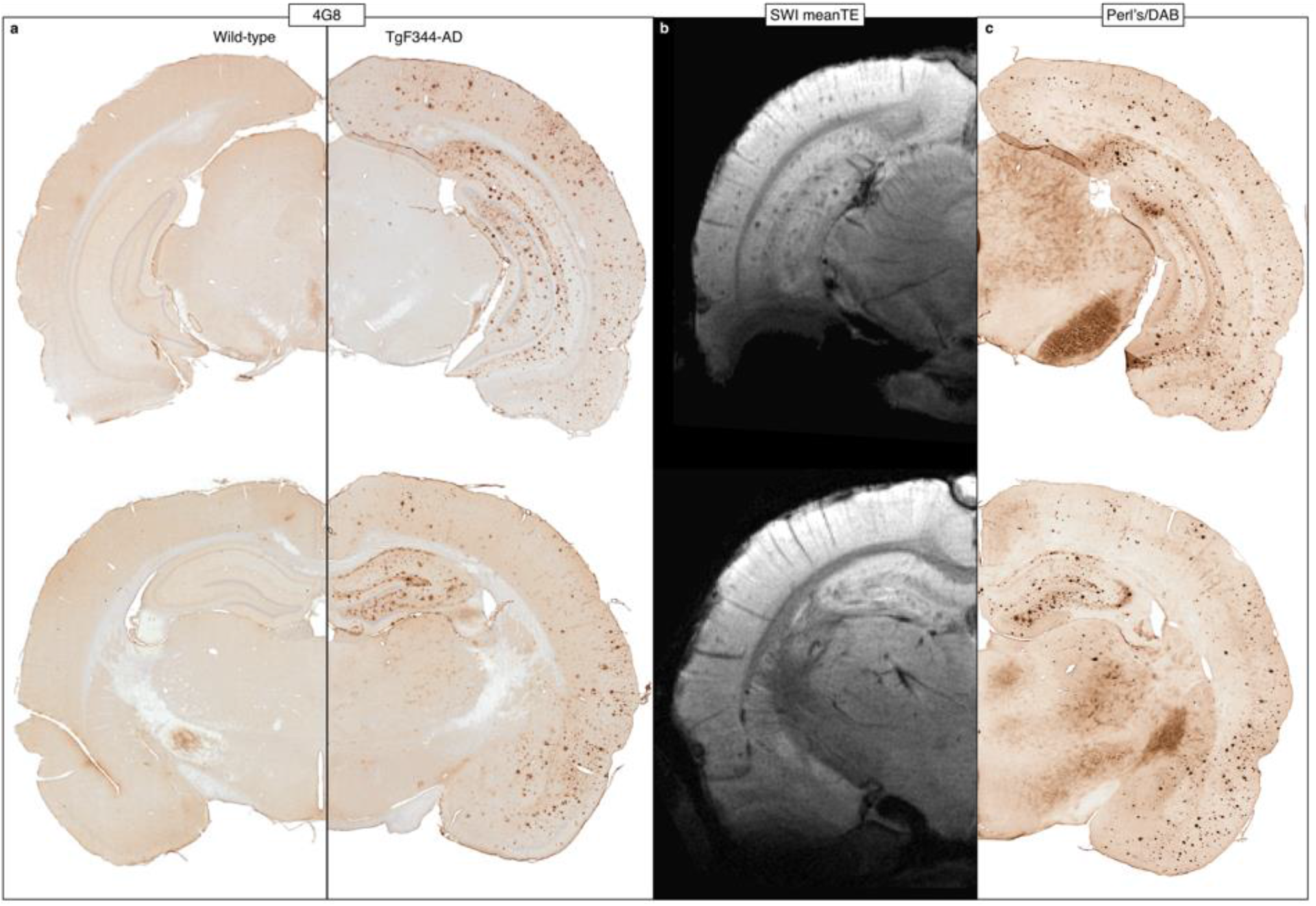
Histology shows iron-containing Aβ plaques throughout the brain in TgF344-AD rats. a) 48G anti-amyloid staining confirmed the presence of plaques throughout the hippocampus and cortex of TgF344-AD rats. b) Hippocampal and cortical plaques were visible on MRI. Susceptibility artifacts around the air-tissue interfaces at the ventral surface of the brain obscured visualization of plaques in those areas. c) Perl’s-DAB staining showed a spatial distribution of iron very similar to Aβ. All images were acquired from one wild-type (a − left) and one TgF344-AD rat (a - right, b, c).

### 3.5 Mn(II) uptake was increased in areas of high plaque burden

The MP2RAGE data revealed a trend (0.22 < FWE-corrected p < 0.5) towards increased R1_norm_ in 5xFAD compared to WT mice in several brain regions including the deep cortical layers, hippocampus, thalamus, and septum (**Figure 11a**,**b**). The spatial pattern of increased R1_norm_ is consistent with the histologically verified pattern of Aβ deposition (**Figure 6a**). This is consistent with the apparent binding of Mn(II) to senile plaques (**Figures 7** and **8**). To illustrate the magnitude of the genotype-driven difference in R1_norm_, an ROI in the anterior cortex was automatically generated by thresholding the voxel-wise statistical map at FWE-corrected p < 0.3 and taking the largest connected component (**Figure 11c**), and the mean R1_norm_ within the ROI was plotted for each mouse (**Figure 11d**). The mean R1_norm_ was significantly greater in 5xFAD mice compared to wild-types (two-sample t-test p = 0.028, Cohen’s *d* = 1.44).

**Figure 11.**
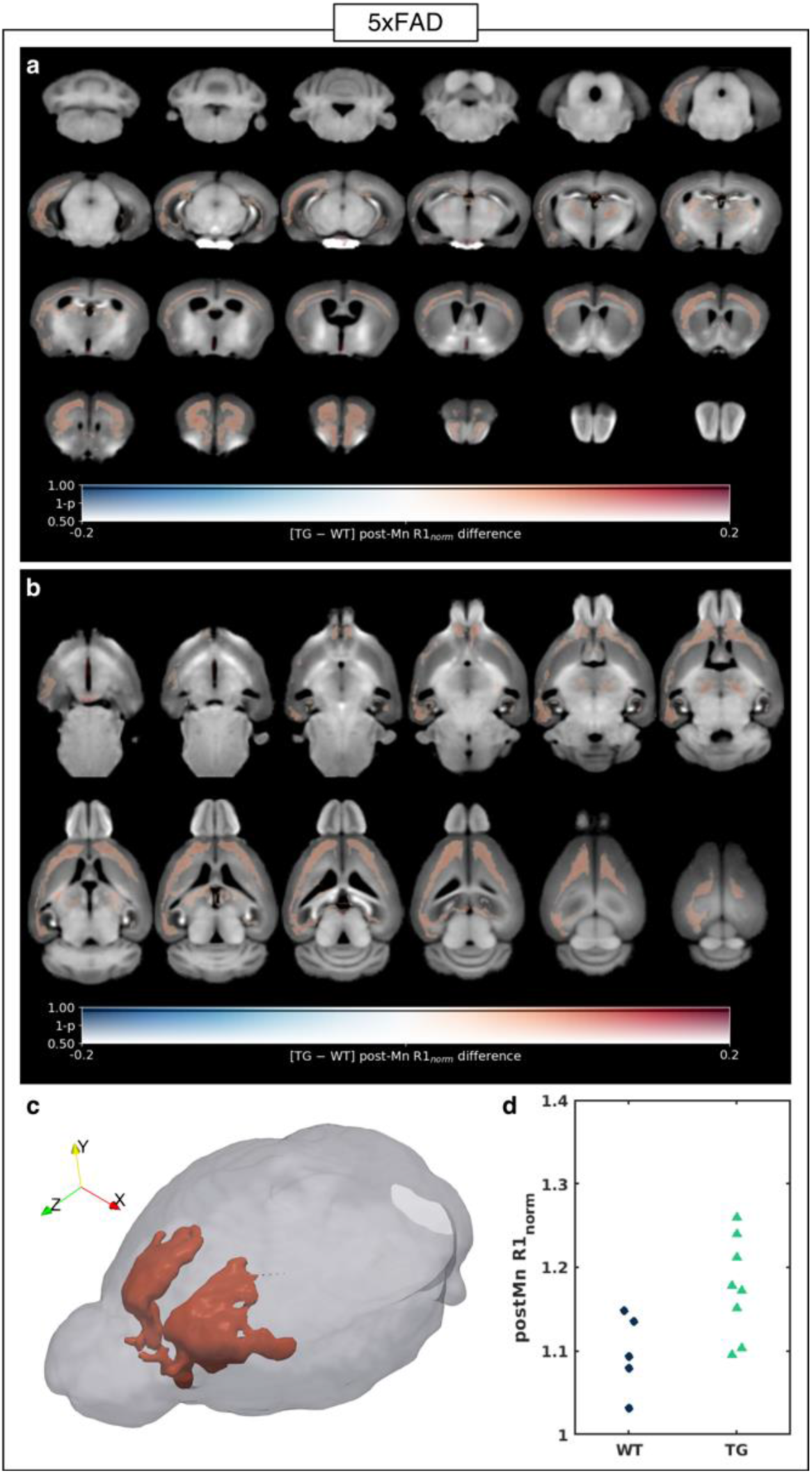
5xFAD mice show increased manganese uptake in the deeper layers of the cortex. a-b) The voxel-wise difference in post-Mn R1_*norm*_ (R1 values were normalized to the median R1 in a manually defined region in the temporalis muscle) between 5xFAD (TG, n=8) mice and wild-type (WT, n=5) littermates, overlaid on the T1-weighted MP2RAGE study-specific template shown in a) coronal slices from back to front and b) transverse slices from bottom to top. The difference in group means is coded by overlay color (warm colors indicate TG > WT), and the statistical significance is coded by overlay transparency (completely transparent indicates family-wise-error-corrected p > 0.5). c) 3D rendering of the template brain (gray) and a cortical region of interest (ROI) automatically generated from the largest connected component in which p < 0.3 (red). d) A dot plot of the mean post-Mn R1_*norm*_ within the ROI for each mouse. Two-tailed two-sample t-test, p = 0.028.

R1_norm_ was also increased in TgF344-AD rats compared to their wild-type littermates but in different areas of the brain than in 5xFAD mice – mostly in the rhinencephalon including the olfactory bulb, amygdala, and piriform cortex (**Figure 12a**,**b**). Also unlike the 5xFAD mice, the increase in R1_norm_ was statistically significant (FWE-corrected p < 0.05) in sizeable clusters (black contours, **Figure 12a**,**b**). A rhinencephalon ROI was automatically segmented by taking the largest connected component of the R1_norm_ p-value map after thresholding it at FWE-corrected p < 0.05 (**Figure 12c**). The mean R1_norm_ in the ROI was significantly greater in TgF344-AD rats than in wild-types (two-sample t-test p < 0.00001, Cohen’s *d* = 12.93, **Figure 12d**).

**Figure 12.**
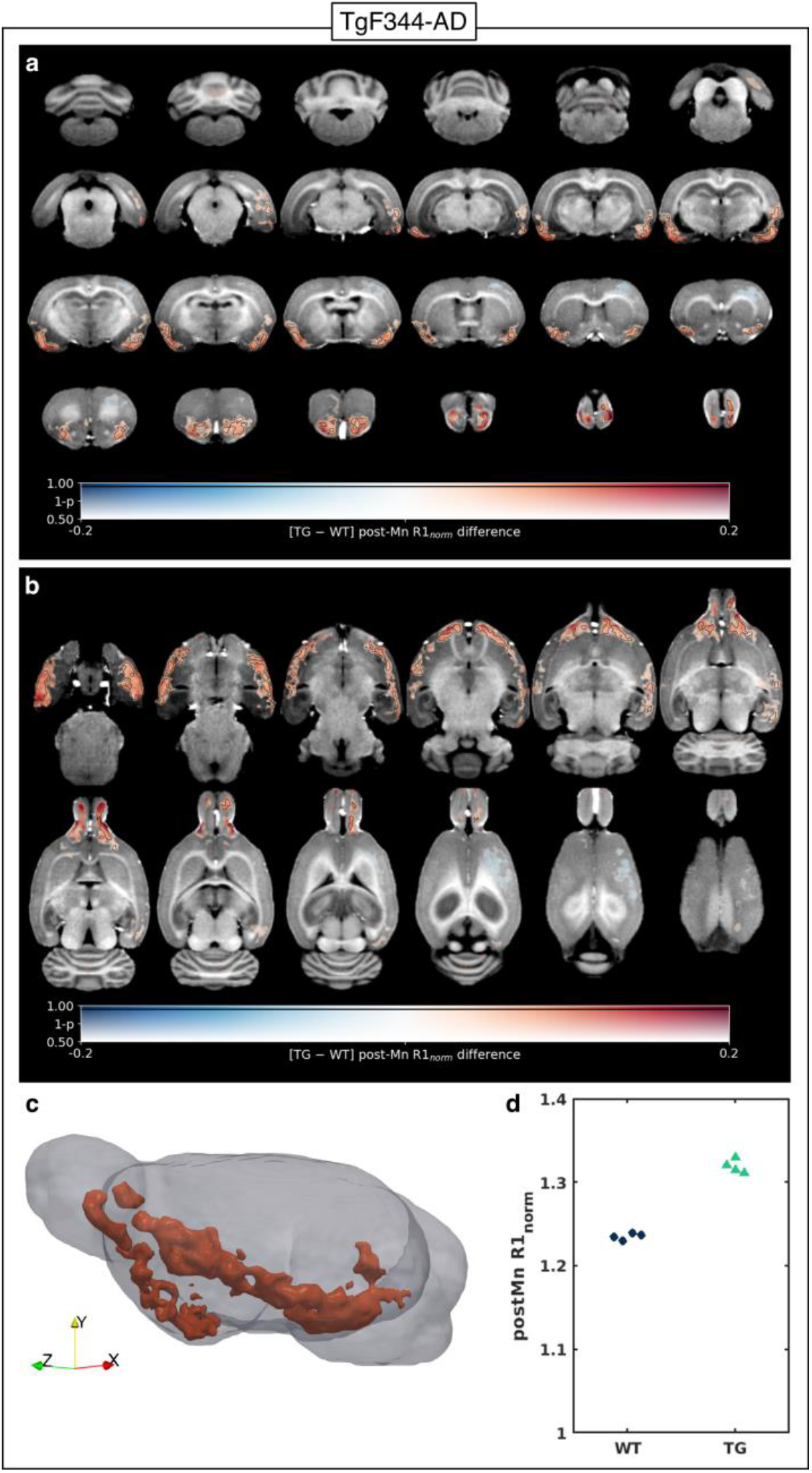
TgF344-AD rats show increased manganese uptake in the rhinencephalon. a-b) The voxel-wise difference in post-Mn R1_*norm*_ (R1 values were normalized to the median R1 in a manually defined region in the temporalis muscle) between TgF344-AD (TG, n=4) rats and wild-type (WT, n=4) littermates, overlaid on the pre-Mn T1-weighted MP2RAGE study-specific template shown in a) coronal slices from back to front and b) transverse slices from bottom to top. The difference in group means is coded by overlay color (warm colors indicate TG > WT), and the statistical significance is coded by overlay transparency (completely transparent indicates family-wise-error-corrected p > 0.5). Areas in which p < 0.05 are outlined in black. c) 3D rendering of the template brain (gray) and a region of interest (ROI) in the rhinencephalon automatically generated from the largest connected component in which p < 0.05 (red). d) A dot plot of the mean post-Mn R1_*norm*_ within the rhinencephalon ROI for each rat. Two-tailed two-sample t-test, p < 0.00001.

## 4 Discussion

### 4.1 MnCl_2_ as a GBCA alternative

The original aim of this study was to evaluate the suitability of MnCl_2_ as an easily deliverable alternative to GBCAs to aid direct visualization of senile plaques in high-resolution MR images. While Mn(II) boosted SNR and tissue contrast, it did not appreciably enhance plaque visibility in 5xFAD mice (**Figure 2**). However, given that the SNR of the pre-Mn images was sufficient to visualize plaques, the signal enhancement provided by Mn(II) could be leveraged to increase the image resolution and/or shorten the scan time while maintaining the ability to detect plaques.

Compared to studies that used GBCAs, we achieved less signal enhancement with MnCl_2_. For example, Santin et al. delivered Gd-DOTA to the brain parenchyma in mice using microbubbles and ultrasound to transiently open the blood-brain barrier, which reduced the cortical T1 from ∼2000 ms to ∼360 ms at 7T (Santin et al., 2013). This dramatic T1 shortening allowed them to acquire images with a resolution of 29×29×117 μm^3^ (∼9× higher than our MGE images) in just 32 minutes. In comparison, we measured a much smaller decrease in whole-brain T1 from ∼1600 ms to ∼1300 ms at 9.4T after MnCl_2_ injections in the rats. This large difference in T1 shortening is due to the much higher Gd-DOTA dose compared to the MnCl_2_ dose given to the rats (4 mmol/kg vs. 0.3 mmol/kg). The toxicity of free manganese prevents the use of such high doses of MnCl_2_, which therefore cannot enable a comparable combination of high resolution and short scan time. Nevertheless, the ease of MnCl_2_ administration by simple subcutaneous injections, compared to complex delivery required for GBCAs is advantageous for many non-invasive applications.

### 4.2 Discrepancy between MRI and histology

There was a discrepancy between the spatial distributions of MR-visible plaques in MGE images and iron-loaded plaques on histological sections. Perl’s staining with potassium ferrocyanide specifically stains Fe(III), but the additional DAB intensification step results in staining of both Fe(III) and Fe(II) (Roschzttardtz et al., 2009). A recent phantom study found that Fe(III) and Fe(II) have significantly different r2* relaxivities: 12.5 mM^-1^s^-1^ and 0.77 mM^-1^s^-1^, respectively (O. Dietrich et al., 2017).

Aβ, like most proteins, is diamagnetic. Putative Aβ plaques have been shown to appear as diamagnetic spots in *ex vivo* QSM of transgenic Aβ mice (Gong et al., 2019). However, many of the plaques in the animal models used in this study appear to contain iron, and the MR-visible plaques have positive susceptibilities. Co-localization of paramagnetic iron with diamagnetic Aβ may reduce susceptibility-based MR contrast.

While Perl’s/DAB staining revealed the presence of plaque-associated iron throughout the brain, regional variation in plaque configuration (compact vs. diffuse) (Dudeffant et al., 2017), relative concentrations of Aβ and iron, and different species of iron might explain why certain plaques were not visible in the MGE images. More nuanced and quantitative molecular analysis is required to test these hypotheses.

Given the relatively large voxel dimensions, the size of the iron core might be the most important determinant of MR visibility, especially in these T2*-weighted MGE images. 4G8 staining shows that plaque sizes do not differ much between the cortex, hippocampus, and thalamus in the 5xFAD mouse (**Figure 6a**,**c**). In contrast, Perl’s/DAB staining shows that the hippocampus contains several iron cores that are much larger than most in the cortex (**Figure 6d**). This could explain why, although there are many more plaques in the cortex, most of the MR-visible plaques are in the hippocampus (**Figure 6b**).

In addition, more plaques were MR-visible in TgF344-AD rats than in 5xFAD mice, supporting the hypothesis that size was the key factor underlying plaque visibility in our MGE images. Qualitatively comparing the two models, plaque size seems to scale proportionally with brain size; and in the TgF344-AD rat, there is no obvious difference in the Aβ to iron ratio within plaques in the cortex compared to those in the hippocampus (**Figures 6** and **10**).

### 4.3 Mn(II) uptake as an indirect marker of senile plaque burden

While MnCl_2_ cannot enable the kind of high-resolution imaging possible with GBCAs, MEMRI did serve another purpose in revealing regional increased Mn(II) uptake and retention in transgenic AD animals compared to wild-types. Previous MEMRI studies have also reported increased Mn(II) in AD mouse models including the 5xFAD model used here, attributing it to neuronal dysfunction and hyperactivity (Fontaine et al., 2017; Tang et al., 2016). This is consistent with the results of Busche et al. who, through *in vivo* measurements of spontaneous Ca^2+^ transients in individual neurons, found hyperactive cortical neurons in the close vicinity (within 60 μm) of Aβ plaques in the APP23xPS45 mouse model of AD (Busche et al., 2008).

We present evidence of another potential mechanism by which Mn(II) retention was enhanced in 5xFAD mice. The increased R1_norm_ in brain regions of high plaque load, in conjunction with Alizarin Red staining of plaques in Mn+ but not Mn-5xFAD mice (**Figure 8**), suggests that accumulation of the injected Mn(II) in these regions was increased by direct binding of Mn(II) to plaques. This is supported by recent studies that showed that Mn(II) binds to Aβ with a weak binding affinity that does not affect the protein’s aggregation (Lermyte et al., 2019; Wallin et al., 2016).

However, Alizarin Red staining of the TgF344-AD rats brains showed that Mn(II) does not have the same affinity to all plaques (**Figure 9**). Moreover, while the plaque load in TgF344-AD rats was equally high in the hippocampus, the increased R1_norm_ was localized to the rhinencephalon. Together, these results indicate that Mn(II) accumulation was increased in the TgF344-AD rat rhinencephalon due to neuronal dysfunction rather than Mn(II) binding to Aβ.

### 4.4 Thalamic calcifications in 5xFAD mice

In addition to senile plaque-like structures in the hippocampal and cortical areas, we also detected large clusters of hypointensities in MGE images in 4/8 5xFAD mice in their bilateral mediodorsal thalami. Similar thalamic lesions have been reported before in transgenic AD mice (Dhenain et al., 2009; Jack et al., 2004), and it has been suggested that these are not typical amyloid plaques, but instead deposits of calcium together with a variable amount of colocalized iron. Accordingly, while we observed a matching pattern between amyloid (4G8) and iron (Perl’s) staining and our MRI-visible plaques in the hippocampus and cortex, neither stain resembled the configuration of the thalamic lesions which were only replicated by the staining for calcium with Alizarin Red (Dahl, 1952). QSM showed that, at pre-Mn baseline, these thalamic clusters were diamagnetic, corroborating the histological findings that these lesions are calcifications.

Interestingly, strikingly similar thalamic calcifications have been observed in animal models of neurotoxicity (Aggarwal et al., 2018; Mori et al., 2000), ischemia/hypoxia (Wideroe et al., 2011), depletion of huntingtin protein (P. Dietrich et al., 2017), tauopathy (Ni et al., 2019), and even ageing (Fraser, 1968). Such dystrophic calcifications (intra or extra-cellular deposits of calcium salts in degenerating or necrotic tissue) are known to occur intracerebrally in the basal ganglia and thalami in a spectrum of human disorders known as primary familial brain calcification (Lemos et al., 2013). A common mechanism that appears to feature in all these calcifications is a disturbed iron homeostasis, as dysregulation of brain transferrin and ferritin has been shown to precede the calcifications. Iron is also known to be pathologically linked to AD and could play a role in the formation of thalamic calcifications in 5xFAD mice.

The regional location of these calcifications could be due to the same reason why calcifications target thalami in the neurotoxicity models; it is possible that an unknown factor related to, e.g., thalamic configuration or accessibility renders it particularly vulnerable. The same could underlie our observation that Mn(II) also appeared to bind to these thalamic deposits, turning them from diamagnetic at pre-Mn baseline to paramagnetic after MnCl_2_ injection (**Figure 7**). This is supported by observations of Mn(II) binding to thalamic calcifications in a MEMRI study of hypoxia induced brain injury (Wideroe et al., 2011), as well as of its binding directly to amyloid plaques (Lermyte et al., 2019; Wallin et al., 2016). Nevertheless, while the observation of Mn-enhancing thalamic calcifications in experimental AD is interesting, the question remains about their wider significance and whether they may be useful biomarkers of either disease progression or treatment efficacy.

### 4.5 Translational outlook

A recent study of MEMRI on healthy volunteers using the FDA-approved, but no longer marketed, mangafodipir demonstrated signal intensity increase in the choroid plexus and anterior pituitary gland but no signal enhancement in the brain parenchyma. This is likely due to the much lower dose (5 μmol/kg) compared to animal studies (0.3 or 0.15 mmol/kg in this study) (Sudarshana et al., 2019). This low dose, necessitated by manganese toxicity, currently limits the clinical translatability of MEMRI for AD applications.

## 5 Conclusion

MnCl_2_ falls short of GBCAs in its ability to enhance longitudinal relaxation, increase image resolution, and reduce scan time. However, it is much simpler to deliver to the brain parenchyma and still offers useful signal enhancement. Our results suggest that, at least in the 5xFAD mouse, this should be leveraged to increase spatial resolution rather than SNR in the context of visualizing senile plaques. Image resolution and plaque size appear to be the key factors in determining plaque visibility, with iron load playing an increasingly important role as resolution and/or plaque size decrease. MEMRI also allows indirect detection of senile plaques. Mn(II) uptake was increased in regions of high plaque burden, consistent with neuronal hyperactivity as a result of plaque-related dysregulation, and perhaps enhanced by Aβ-Mn(II) binding. Thus, MEMRI is a viable method for visualizing senile plaques and for obtaining functional insights in preclinical models of AD. This technique will be used in future longitudinal studies to monitor disease progression and therapeutic response.

## 6 Acknowledgements

Funding: This work was supported by the Alzheimer’s Society [AS-PhD-18B-015] and internal funding as “pump-priming” for developing novel imaging methodologies.

The authors thank the Wohl Cellular Imaging Centre (http://www.kclwcic.co.uk) for the use of their slide scanner. SW would also like to thank the Wellcome Trust and Medical Research Council for their ongoing support of our neuroimaging research.

Declarations of interest: none

## Abbreviations

AD: Alzheimer’s disease
Aβ: beta-amyloid
CNR: contrast-to-noise ratio
DAB: 3,3′-Diaminobenzidine
FWE: family-wise error
GBCA: gadolinium-based contrast agents
MRI: magnetic resonance imaging
MEMRI: manganese-enhanced MRI
MGE: multi-gradient-echo
Mn-: MnCl_2_-naïve
Mn+: treated with MnCl_2_
MP2RAGE: magnetization prepared 2 rapid acquisition of gradient echoes
PBS: phosphate-buffered saline
QSM: quantitative susceptibility mapping
ROI: region-of-interest
SNR: signal-to-noise ratio
SWI: susceptibility-weighted image
TBS: Tris-buffered saline
TE: echo time
TI: inversion time
UTE: ultra-short echo time

## References

Aggarwal, M., et al. (2018). Nuclei-specific deposits of iron and calcium in the rat thalamus after status epilepticus revealed with quantitative susceptibility mapping (QSM). J Magn Reson Imaging, 47(2), 554–564. doi:10.1002/jmri.25777

Avants, B. B., et al. (2008). Symmetric diffeomorphic image registration with cross-correlation: evaluating automated labeling of elderly and neurodegenerative brain. Med Image Anal, 12(1), 26–41. doi:10.1016/j.media.2007.06.004

Badea, A., et al. (2019). Multivariate MR biomarkers better predict cognitive dysfunction in mouse models of Alzheimer’s disease. Magn Reson Imaging, 60, 52–67. doi:10.1016/j.mri.2019.03.022

Brandt, M., et al. (2019). Manganese in PET imaging: Opportunities and challenges. J Labelled Comp Radiopharm, 62(8), 541–551. doi:10.1002/jlcr.3754

Busche, M. A., et al. (2008). Clusters of hyperactive neurons near amyloid plaques in a mouse model of Alzheimer’s disease. Science, 321(5896), 1686–1689. doi:10.1126/science.1162844

Cohen, R. M., et al. (2013). A transgenic Alzheimer rat with plaques, tau pathology, behavioral impairment, oligomeric abeta, and frank neuronal loss. J Neurosci, 33(15), 6245–6256. doi:10.1523/JNEUROSCI.3672-12.2013

Dahl, L. K. (1952). A simple and sensitive histochemical method for calcium. Proc Soc Exp Biol Med, 80(3), 474–479. doi:10.3181/00379727-80-19661

Dhenain, M., et al. (2009). Characterization of in vivo MRI detectable thalamic amyloid plaques from APP/PS1 mice. Neurobiol Aging, 30(1), 41–53. doi:10.1016/j.neurobiolaging.2007.05.018

Dietrich, O., et al. (2017). MR imaging differentiation of Fe(2+) and Fe(3+) based on relaxation and magnetic susceptibility properties. Neuroradiology, 59(4), 403–409. doi:10.1007/s00234-017-1813-3

Dietrich, P., et al. (2017). Elimination of huntingtin in the adult mouse leads to progressive behavioral deficits, bilateral thalamic calcification, and altered brain iron homeostasis. PLoS Genet, 13(7), e1006846. doi:10.1371/journal.pgen.1006846

Dudeffant, C., et al. (2017). Contrast-enhanced MR microscopy of amyloid plaques in five mouse models of amyloidosis and in human Alzheimer’s disease brains. Sci Rep, 7(1), 4955. doi:10.1038/s41598-017-05285-1

Fontaine, S. N., et al. (2017). Identification of changes in neuronal function as a consequence of aging and tauopathic neurodegeneration using a novel and sensitive magnetic resonance imaging approach. Neurobiol Aging, 56, 78–86. doi:10.1016/j.neurobiolaging.2017.04.007

Fraser, H. (1968). Bilateral thalamic calcification in ageing mice. J Pathol Bacteriol, 96(1), 220–222. doi:10.1002/path.1700960124

Gong, N. J., et al. (2019). Imaging beta amyloid aggregation and iron accumulation in Alzheimer’s disease using quantitative susceptibility mapping MRI. Neuroimage, 191, 176–185. doi:10.1016/j.neuroimage.2019.02.019

Helms, G., & Dechent, P. (2009). Increased SNR and reduced distortions by averaging multiple gradient echo signals in 3D FLASH imaging of the human brain at 3T. J Magn Reson Imaging, 29(1), 198–204. doi:10.1002/jmri.21629

Jack, C. R., Jr., et al. (2018). NIA-AA Research Framework: Toward a biological definition of Alzheimer’s disease. Alzheimers Dement, 14(4), 535–562. doi:10.1016/j.jalz.2018.02.018

Jack, C. R. Jr.,, et al. (2004). In vivo visualization of Alzheimer’s amyloid plaques by magnetic resonance imaging in transgenic mice without a contrast agent. Magn Reson Med, 52(6), 1263–1271. doi:10.1002/mrm.20266

Lemos, R. R., et al. (2013). An update on primary familial brain calcification. Int Rev Neurobiol, 110, 349–371. doi:10.1016/B978-0-12-410502-7.00015-6

Lermyte, F., et al. (2019). Metal Ion Binding to the Amyloid beta Monomer Studied by Native Top-Down FTICR Mass Spectrometry. J Am Soc Mass Spectrom, 30(10), 2123–2134. doi:10.1007/s13361-019-02283-7

Massaad, C. A., & Pautler, R. G. (2011). Manganese-enhanced magnetic resonance imaging (MEMRI). Methods Mol Biol, 711, 145–174. doi:10.1007/978-1-61737-992-5_7

Mathis, C. A., et al. (2012). Development of positron emission tomography beta-amyloid plaque imaging agents. Semin Nucl Med, 42(6), 423–432. doi:10.1053/j.semnuclmed.2012.07.001

Mori, F., et al. (2000). Widespread calcium deposits, as detected using the alizarin red S technique, in the nervous system of rats treated with dimethyl mercury. Neuropathology, 20(3), 210–215. doi:10.1046/j.1440-1789.2000.00341.x

Ni, R., et al. (2019). Tau deposition is associated with imaging patterns of tissue calcification in the P301L mouse model of human tauopathy. bioRxiv, 851915. doi:10.1101/851915

O’Brien, K. R., et al. (2014). Robust T1-weighted structural brain imaging and morphometry at 7T using MP2RAGE. PLoS One, 9(6), e99676. doi:10.1371/journal.pone.0099676

Oakley, H., et al. (2006). Intraneuronal beta-amyloid aggregates, neurodegeneration, and neuron loss in transgenic mice with five familial Alzheimer’s disease mutations: potential factors in amyloid plaque formation. J Neurosci, 26(40), 10129–10140. doi:10.1523/JNEUROSCI.1202-06.2006

Oguz, I., et al. (2014). RATS: Rapid Automatic Tissue Segmentation in rodent brain MRI. J Neurosci Methods, 221, 175–182. doi:10.1016/j.jneumeth.2013.09.021

Perez, P. D., et al. (2013). In vivo functional brain mapping in a conditional mouse model of human tauopathy (tauP301L) reveals reduced neural activity in memory formation structures. Mol Neurodegener, 8, 9. doi:10.1186/1750-1326-8-9

Petiet, A., et al. (2012). Gadolinium-staining reveals amyloid plaques in the brain of Alzheimer’s transgenic mice. Neurobiol Aging, 33(8), 1533–1544. doi:10.1016/j.neurobiolaging.2011.03.009

Robinson, S. D., et al. (2017). Combining phase images from array coils using a short echo time reference scan (COMPOSER). Magn Reson Med, 77(1), 318–327. doi:10.1002/mrm.26093

Roschzttardtz, H., et al. (2009). Identification of the endodermal vacuole as the iron storage compartment in the Arabidopsis embryo. Plant Physiol, 151(3), 1329–1338. doi:10.1104/pp.109.144444

Saar, G., & Koretsky, A. P. (2018). Manganese Enhanced MRI for Use in Studying Neurodegenerative Diseases. Front Neural Circuits, 12, 114. doi:10.3389/fncir.2018.00114

Santin, M. D., et al. (2013). Fast in vivo imaging of amyloid plaques using mu-MRI Gd-staining combined with ultrasound-induced blood-brain barrier opening. Neuroimage, 79, 288–294. doi:10.1016/j.neuroimage.2013.04.106

Schindelin, J., et al. (2012). Fiji: an open-source platform for biological-image analysis. Nat Methods, 9(7), 676–682. doi:10.1038/nmeth.2019

Sudarshana, D. M., et al. (2019). Manganese-Enhanced MRI of the Brain in Healthy Volunteers. AJNR Am J Neuroradiol, 40(8), 1309–1316. doi:10.3174/ajnr.A6152

Tang, X., et al. (2016). Spatial learning and memory impairments are associated with increased neuronal activity in 5XFAD mouse as measured by manganese-enhanced magnetic resonance imaging. Oncotarget, 7(36), 57556–57570. doi:10.18632/oncotarget.11353

Ten Kate, M., et al. (2018). MRI predictors of amyloid pathology: results from the EMIF-AD Multimodal Biomarker Discovery study. Alzheimers Res Ther, 10(1), 100. doi:10.1186/s13195-018-0428-1

Tustison, N. J., et al. (2010). N4ITK: improved N3 bias correction. IEEE Trans Med Imaging, 29(6), 1310–1320. doi:10.1109/TMI.2010.2046908

Wallin, C., et al. (2016). Characterization of Mn(II) ion binding to the amyloid-beta peptide in Alzheimer’s disease. J Trace Elem Med Biol, 38, 183–193. doi:10.1016/j.jtemb.2016.03.009

Wei, H., et al. (2015). Streaking artifact reduction for quantitative susceptibility mapping of sources with large dynamic range. NMR Biomed, 28(10), 1294–1303. doi:10.1002/nbm.3383

Wideroe, M., et al. (2011). Longitudinal manganese-enhanced magnetic resonance imaging of delayed brain damage after hypoxic-ischemic injury in the neonatal rat. Neonatology, 100(4), 363–372. doi:10.1159/000328705

Winkler, A. M., et al. (2014). Permutation inference for the general linear model. Neuroimage, 92, 381–397. doi:10.1016/j.neuroimage.2014.01.060

Wood, T. C. (2018). QUIT: QUantitative Imaging Tools. Journal of Open Source Software, 3(26), 656. doi:10.21105/joss.00656

